# Toward a generalizable deep CNN for neural drive estimation across muscles and participants

**DOI:** 10.1101/2022.08.31.505855

**Authors:** Yue Wen, Sangjoon J. Kim, Simon Avrillon, Jackson T. Levine, François Hug, José L. Pons

## Abstract

High-density electromyography (HD-EMG) decomposition algorithms are used to identify individual motor unit spike trains, which collectively constitute the neural code of movements, to predict motor intent. This approach has advanced from offline to online decomposition, from isometric to dynamic contractions, leading to a wide range of neural-machine interface applications. However, current online methods need offline retraining when applied to the same muscle on a different day or to a different person, which limits their applications in a real-time neural-machine interface. We proposed a deep convolutional neural network (CNN) framework for neural drive estimation, which captures general spatiotemporal properties of motor unit action potentials to generalize its application without retraining to HD-EMG data recorded in separate sessions, muscles, and participants. We recorded HD-EMG signals from the vastus medialis and vastus lateralis muscles while participants performed isometric contractions during two sessions separated by approximately 20 months. We identified motor unit spike trains from HD-EMG signals using a blind source separation (BSS) method, and then used the cumulative spike train (CST) of these motor units and the HD-EMG signals to train and validate the deep CNN. On average, the correlation coefficients between CST from BSS and that from deep CNN were 0.977±0.007 for leave-one-out across-sessions-and-muscles validation and 0.985±0.005 for leave-one-out across-participants validation. When trained with more than four datasets, the performance of deep CNN saturated at 0.979±0.001 for cross validations across muscles, sessions, and participants. Therefore, we can conclude that the deep CNN is generalizable across the afore-mentioned conditions without retraining. We could potentially generate a robust deep CNN to estimate neural drive to muscles for neural-machine interfaces.

## 1. Introduction

Surface electromyography (EMG) has wide applications for neural-machine interfacing because of its non-invasive access to the neural drive to muscles, i.e., the net output of all the motor neurons that innervate the muscle. Various control methods, including pattern recognition, regression models, and musculoskeletal models, have been used to control wearable robots (e.g., prostheses and exoskeleton) using surface EMG [1–4]. Essentially, the neural drive to muscles is composed of discrete spike signals, which are then amplified and transformed to continuous motor unit action potential (MUAP) signals by the innervated muscle. When measured at the surface of the skin, the MUAP signals from all activated MUs are overlapped and temporal-spatially filtered by the tissues between muscle fibers and the skin. Therefore, surface EMG is a rough estimation and is not linearly related to the neural drive to muscles due to the superimposition and cancellation of MUAPs [5].

To have a more accurate estimation of the neural drive to muscles, several motor unit identification techniques have been developed, allowing the decomposition of high-density surface EMG (HD-EMG) into MU spike trains. Holobar et al. combined a Convolution Kernel Compensation (CKC) method and a gradient descent algorithm to extract multiple MU spike trains simultaneously [6, 7]. Alternatively, Chen et al. proposed a progressive FastICA peel-off method for fast and accurate HD-EMG decomposition [8]. Both methods were validated through a two-source validation protocol [6, 9], and they demonstrated a high agreement in MU identification [10]. Recently, Negro et al. proposed a general framework for MU identification using multi-channel invasive and non-invasive EMG signals by combining FastICA and CKC algorithms [11]. The feasibility of these methods for HD-EMG decomposition has been extensively validated with different muscles (from upper limb and lower limb), with different clinical populations [12–14], and for different purposes (pathology studies, interfacing with assistive devices).

Recently, the neural drive to muscles has been used to extract motor intent for human-machine interfaces. Joint torque was estimated using the cumulative spike train (CST) [15], a motor unit twitch model [16], and a MU-specific image-based deep convolutional neural network (CNN) [17]. Joint kinematics were also estimated using MU discharge times [18] with forward kinematics [19], the first principal component of MU firing rates [20], and the CST [21]. Furthermore, these studies demonstrated the improved reliability and accuracy of force and kinematic estimations over traditional EMG-amplitude based approaches. To apply these estimations and achieve real-time human-machine interfacing, efforts have been made to push MU identification to an online platform by applying separation matrices, initialized offline, to real-time HD-EMG data [22, 23]. Additionally, MU identification of dynamic movements has recently been achieved during tasks such as flexion and extension of the elbow and gait [24–26]. Both online and dynamic methods were based on template matching of MUAP, using features stored in a separation matrix that were periodically updated to account for MUAP changes due to muscle fatigue and changes in muscle length.

Although the periodical update strategy could account for slow and gradual MUAP changes, the separation matrix approach is limited to a fixed number of MUs after offline initialization, which restricts its application to the same muscle within the same session. The number of recruited MUs during a contraction is different for different muscles and different participants, and the amplitude and duration of the MUAP are affected by the diameter of the muscle fibers [27, 28]. Therefore, the MUAP shapes are also different across muscles and participants. In addition, when HD-EMG signals are recorded on the surface of the skin, all MUAPs features are influenced by the relative position between the muscle fiber and the electrodes, as well as the tissue in between [29]. Moreover, the MUAP shapes from the same muscle could also change when recorded from a different day or session. Therefore, the separation matrix approach always requires offline initialization for each application to different muscles, different participants, or the same muscle in a different session. A generalizable approach for neural drive estimation across the aforementioned conditions would dramatically reduce the initialization efforts and simplify real-time human-robot interfacing.

The purpose of this study is to validate the generalizability of a deep CNN framework for neural drive estimation across sessions, muscles, and participants. A deep CNN is chosen because it demonstrated success 1) in many complex image pattern recognition problems regardless of the orientation and distortion of the objects [30, 31]; 2) in EMG and electroencephalography analysis as well as MU identification using HD-EMG [32, 33]. In addition, we have demonstrated that the deep CNN is feasible for neural drive estimation across different contraction intensities and joint angles in our previous study [34]. As the deep CNN is a non-linear complex feature extraction approach, it could handle more challenging situations and store much more information than a linear separation matrix. In this study, we first identified the MU spike trains from HD-EMG signals using blind source separation (BSS) and then used both HD-EMG and corresponding CST to train the deep CNN. We validated the generalizability of the deep CNN for within participant variations (i.e., across muscle-session variations), across participants variations, and across muscle-session-participant variations.

## 2. Method

### 2.1. Experiment

#### 2.1.1. Participants

Five healthy participants were enrolled in this study (30±6 years, height 183±6 cm, body mass 74±8 kg). All participants had no history of lower leg injury within the past 6 months. The institutional research ethics committee “Comité de protection des personnes Ile de France XI” approved this study (CPP-MIP-013), and all procedures were in accordance with the Declaration of Helsinki. All participants provided their written informed consent prior to participation in the study. Note that the experimental data were a subset of a previously published dataset [35].

#### 2.1.2. Experimental protocol

The experimental protocol included two experiment sessions (i.e., S1 and S2) separated by ∼ 20 months (± 1 month). During each session, participants were seated on a dynamometer (Biodex System 3 Pro, Biodex Medical, Shirley, NY) with their torso immobilized with two inextensible straps that crossed the torso. Their hip joint was positioned at 90°, where 0° is the neutral position; their knee joint was positioned at 80°, where 0° is full extension (Fig. 1). After a series of warm-up contractions, participants performed three maximal voluntary (isometric) contractions (MVCs) for 3-5 s with two minutes of rest in between. Then, participants performed three submaximal isometric contractions by tracking a target displayed on a screen corresponding to 25% of their peak torque. The target followed a trapezoidal trajectory with a 10-s linear ramp-up, a 10-s plateau, and a 10-s linear ramp-down phase. Each contraction was followed by 30 s of rest. Similarly, on the second visit, participants performed three trapezoidal contractions at 25% of their peak torque with a 5-s linear ramp-up, a 20-s plateau, and a 5-s ramp-down phase. Each contraction was separated by a 60-s period of rest. Torque signals were collected using the same Quattrocento system used for HD-EMG acquisition (in the following Section)

**Figure 1.**
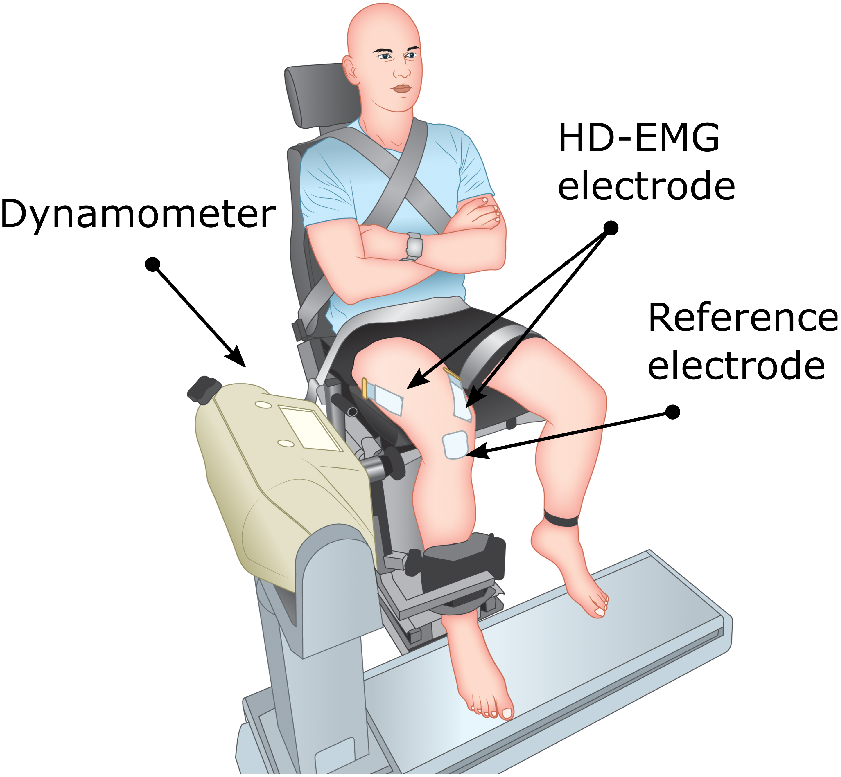
Experimental setup. Participants were seated on a dynamometer (Biodex System 3 Pro, Biodex Medical, Shirley, NY) with their knee joint positioned at 80°. HD-EMG signals were recorded from the vastus lateralis (VL) and the vastus medialis (VM) muscles while participants performed isometric contractions.

#### 2.1.3. Data acquisitio

During all contraction trials, HD-EMG signals were recorded from the vastus lateralis (VL) and the vastus medialis (VM). Two-dimensional adhesive grids of 64 electrodes [13×5 gold-coated electrodes with one electrode absent on a corner; interelectrode distance: 8 mm (ELSCH064NM2, OT Bioelettronica, Italy)] were placed over the VL and VM muscles aligned parallel to the direction of the muscle fascicles. Prior to placing the grids, the skin was shaved and cleaned with an abrasive pad and alcohol. Semi-disposable bi-adhesive foam layers (SpesMedica, Battipaglia, Italy) were used to attach the adhesive grids on the skin. These foam layers were equipped with cavities such that conductive paste (SpesMedica, Battipaglia, Italy) could fill the cavities to make skin-electrode contact. Reference electrodes (Kendall Medi-Trace, Canada) were placed over the patella. The multi-channel acquisition system (EMG-Quattrocento; 400-channel EMG amplifier, OT Bioelettronica, Italy) was used to record EMG signals in monopolar mode, which was then band-pass filtered (10-900 Hz) and digitized at a sampling rate of 2048 Hz.

#### 2.1.4. HD-EMG decompositio

The HD-EMG signals from all channels were band-pass filtered offline (Butterworth 2nd order, 20–500 Hz) and were visually inspected to remove noisy channels with low signal-to-noise ratio or motion artifacts. After pre-processing the data, the HD-EMG signals were decomposed into MU spike trains using a convolution BSS method [11], which has been extensively validated using experimental and simulated signals. Then, all MU spike trains were visually inspected and manually edited to remove any false positives (FP; labeled artifact) or false negatives (FN; non-labeled discharges) by an experienced operator. Only the MUs that exhibited a pulse-to-noise ratio (PNR) greater than 30 dB were retained for further analysis. This threshold ensured a sensitivity higher than 90% and a FP rate lower than 2% [29].

### 2.2. Neural drive estimation using deep CNN

#### 2.2.1. Deep CNN for neural drive estimation

We have previously developed a deep CNN framework to directly estimate the neural drive in the form of the CST [34]. The deep CNN identified the number of MU spikes in a given window of HD-EMG signals. The deep CNN took the window of HD-EMG signals as its input where the width of the window was M (the number of HD-EMG channels) and the length of the window was W (e.g., 40 data points from each channel when window size equals 40). A sliding window approach, which is commonly used in offline EMG analysis methods [15,23,36], was used to segment the data with the increment of the window defined as the step size. Then the deep CNN estimated the number of MU spikes within a given window as its output, and if no spikes was identified, the output was set to zero for all nodes (see Fig. 2). A node was set equal to one for every spike identified within a window (e.g., for two spikes identified, the first two nodes were set to one and the rest to zero).

**Figure 2.**
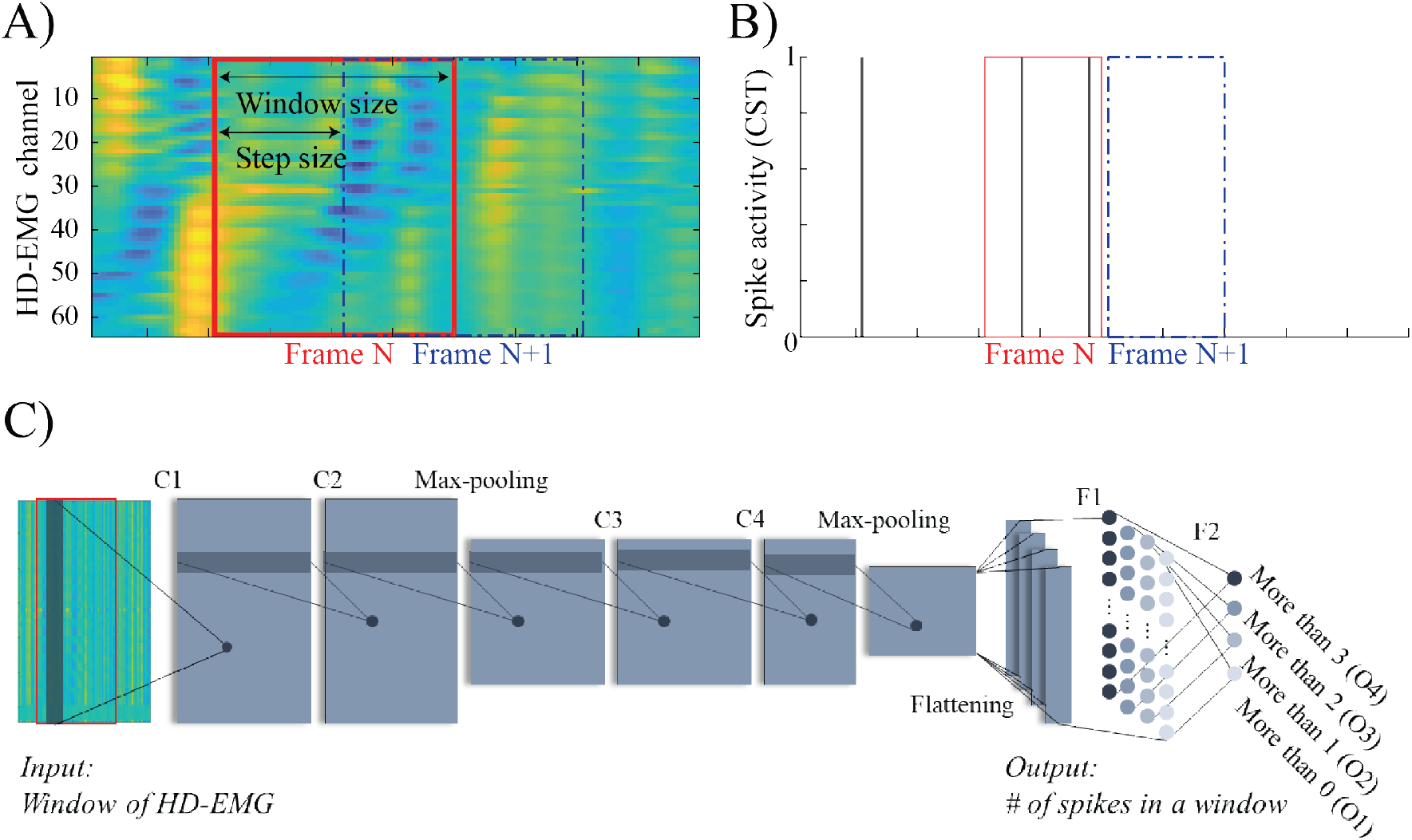
A) The input of the deep CNN: the HD-EMG signals were segmented into windows using a sliding window method with a window size of 40 and a step size of 20, generating an overlap of 20 data points between windows. B) The output of the deep CNN: the CST decomposed using BSS was also segmented into windows using a similar sliding window method with a window size of 20 and a step size of 20, generating no overlap between windows to avoid double counting. C) Deep CNN framework for neural drive estimation in form of CST.

The full structural design of the deep CNN is presented in [34]. In short, the deep CNN consists of six parametric layers (four convolutional layers and two fully connected layers) and three non-parametric layers (two max pooling layers and one flatten layer) between the input and output layers. A sigmoid activation function, ranging from 0 to 1, was used for the output layer and a rectified linear unit (ReLU) activation function was used for all other layers. Additionally, the consecutive convolutional layers (C1 and C2, C3 and C4) were designed such that the model would be capable of learning complex features while the fully connected layers were designed to make predictions based on feature maps that the convolutional layers provided. Previously, we have demonstrated the ability to generalize this deep CNN across different contraction intensities, contraction target profiles (e.g., trapezoidal vs sinusoidal), and muscle geometries (e.g., for different joint angles). Here we aim to demonstrate the ability to generalize across sessions and muscles, and across participants.

#### 2.2.2 Data preparatio

The HD-EMG signals were visually inspected, and channels with low signal-to-noise ratio or motion artifacts were assigned the mean values of the signals from surrounding channels. Additionally, we aligned the spike indices to the center of the MUAP. First, the MUAP shape was extracted using the spike indices of each MU. Then, spike indices were adjusted to align with the center of the MUAP. After pre-processing, we used the same sliding window method as in [34] to segment the HD-EMG signals and CST into small windows. For the HD-EMG windowing, we used a window length of 40 data points and sliding step of 20 data points with an overlap of 20 data points (Fig. 2A). For the CST windowing, we used a window length of 20 data points and sliding step of 20 data points without overlapping to avoid double counting (Fig. 2B). The CST frame and HD-EMG frame were aligned in the center. The summation of the spikes in each CST frame was used to label the corresponding HD-EMG frame. Based on our previous study, we limited the number of outputs of the deep CNN to 4. If there was one spike in the window, the first node was set to one; if there were two spikes in the window, the first two nodes were set to 1. However, if there were more than 4 spikes in the window, we set all four nodes to 1. Therefore, we might lose some of the spikes in the training and testing data, but this is the best choice when comparing the output of the deep CNN with the full CST extracted from the BSS [34].

#### 2.2.3. Training and validation of deep CNN

For within-participant validation, we validated the generalizability of the deep CNN in estimating the neural drive across two factors (i.e., muscle and session) using four datasets (i.e., S1L and S1M from session one, S2L and S2M from session two; L refers to VL muscle, and M refers to VM muscle) from each participant. Since the performance of the deep CNN heavily relies on the training and validation datasets, we organized the data into two formats: First, we grouped data from three contractions of each muscle-session condition to obtain one fold of data; Second, we grouped each contraction from four muscle-session conditions to create one fold of data (Fig. 3A). Accordingly, we performed two types of cross-validations: 1) a standard 3-fold cross-validation (CV), where each fold included data from one contraction from each dataset (Fig. 3B) and two folds were used for training and one fold was used for validation; 2) a leave-one-out cross-validation (LOOCV), where three datasets were used for training and one dataset was left out for validation (Fig. 3C); The 3-fold CV is to validate the capacity of the deep CNN when it has full access to the datasets. The LOOCV is to validate the performance of the deep CNN when trained and tested with different datasets. We performed two types of cross-validations for each participant.

**Figure 3.**
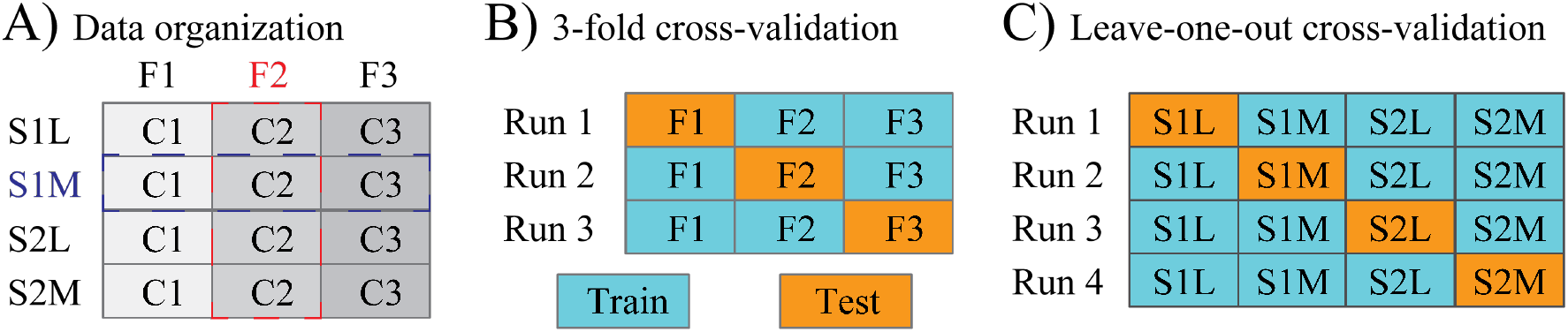
Data organization and validation methods: A) Data organization. The experimental data was organized in two ways: 1) Fold 1 to Fold 3 represented the three contractions of each dataset (S1L, S1M, S2L, and S2M); 2) S1L, S1M, S2L and S2M represented the four dataset from two muscles in two separated sessions; S1 and S2 represent the first and second session, respectively; L and M represent the VL and VM muscles, respectively. B) The standard 3-fold cross-validation (3CV). For ‘Run 1’, two folds (F2 and F3) were used for training and one fold (F1) was used for validation. Three runs were performed to get the averaged performance. C) The leave-one-out cross-validation (LOOCV). For ‘Run 1’, three datasets (S1M, S2L, and S2M) were used for training and one dataset (S1L) was left out for validation. Four runs were performed to get the averaged performance.

For across-participant validation, we also performed 3-fold CV and LOOCV to investigate the generalizability of the deep CNN for neural drive estimation across participants using 5 datasets (one from each participant). For the 3-fold CV, each fold included data from five contractions, one from each participant; two folds were used to train the deep CNN and the remaining fold was used to validate the deep CNN. For LOOCV validation, the deep CNN was trained with data from four participants and validated with data from the left-out participant. We performed the aforementioned two types of cross-validations for each muscle in each session.

In addition, we validated the generalizability of the deep CNN across all conditions regardless of the muscles, sessions, and participants. Among 20 datasets (5P*2S*2M), we randomly selected a subset of data for training and used the remaining datasets for testing. Here we also investigated how the training data size (i.e., the number of datasets included for training) affected the generalizability of the deep CNN. First, we trained the deep CNN with each one of the 20 datasets and tested with the remaining 19 datasets. Here, we ran 20 iterations for all possible combinations as a baseline performance. Then, we trained the deep CNN with combinations of multiple datasets (up to 6 datasets) and tested with the remaining datasets. For each training data size condition, we ran 5 iterations, each with a randomly selected fixed number of datasets for training, to get representative results.

### 2.3. Performance evaluation

The outputs of the deep CNN were summed to form the CST, which was then smoothed using a 400-ms Hanning window to estimate the neural drive [37]. We calculated the correlation coefficient and the normalized root-mean-square error (nRMSE) between the smoothed CST from manually edited MU spike trains from BSS and that from the deep CNN to evaluate the accuracy of deep CNN in neural drive estimation. Specifically, the nRMSE was defined as

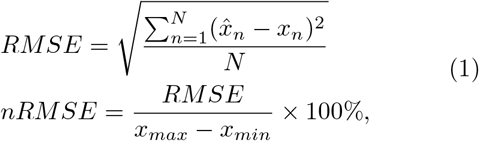

where 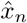 and *x*_*n*_ are the nth sample of the smoothed CST from deep CNN and the smoothed CST from BSS, respectively. *x*_*max*_ and *x*_*min*_ are the maximum and minimum values of the smoothed CST from BSS, and N is the number of samples.

The deep CNN could generate false positive errors during the neural drive estimation, i.e., identifying spike activities while the muscle is not activated. Therefore, we included 2-s data before and after the actual muscle contraction when calculating the correlation coefficient and nRMSE.

### 2.4. Statistical Analyses

A two-factor ANOVA (factor 1: validation methods; factor 2: data sources) was performed to test whether the performance of the deep CNN was validation method dependent or data source dependent. Tukey’s honestly significant difference procedure was used to test the statistical significance between conditions. Here, data sources refer to datasets from each one of the five participants for the within-participant validation and datasets from each one of the four session-muscle combinations for across-participant validation. On one hand, if the deep CNN performed well in 3-fold CV method but significantly worse in LOOCV method, this could indicate that the knowledge might not be transferable from one participant to another participant or from one muscle to another muscle. On the other hand, if the deep CNN performed well with data from one participant/muscle but much worse with data from another participant/muscle, this could indicate that the data quality has a significant effect on the performance of the deep CNN. Additionally, a Pearson correlation test was performed to quantify the relationship between the generalizability of the deep CNN and the number of MUs to determine the Pearson correlation coefficient (r) and its associated P value (*α* = .05). All analyses were conducted using custom-written scripts in Matlab 2020.

## 3. Results

### 3.1. Number of MUs and spikes identified using BSS

The number of MUs and total number of spikes from five participants across four experimental conditions are reported in Table. 1. The number of MUs and the number of spikes varied across experimental conditions (Table. 1). For session 1, across participants, the average number of MUs identified using BSS was 28±7 for VL muscle and 16±4 for VM muscle; the number of spikes was 15700±6229 for VL muscle and 8808±3395 for VM muscle. For session 2, the number of MUs identified using BSS was 34±3 for VL muscle and 15±2 for VM muscle; the number of spikes was 21400±2794 for VL muscle and 10023±2053 for VM muscle. Similarly, the number of MUs and the number of spikes varied across participants (Table. 1). Averaged across all four experimental conditions, P2 was the highest in both the number of MUs (26±9) and the number of spikes (15279±5449), while P3 was the lowest in both the number of MUs (20±13) and the number of spikes (10624±8704).

**Table 1.** The number of MUs and number of spikes for all participants and experimental conditions.

|  | Session 1 |  |  |  | Session 2 |  |  |  |
| --- | --- | --- | --- | --- | --- | --- | --- | --- |
|  | VL |  | VM |  | VL |  | VM |  |
|  | # of MUs | # of spikes | # of MUs | # of spikes | # of MUs | # of spikes | # of MUs | # of spikes |
| P1 | 36 | 18290 | 20 | 8827 | 30 | 17581 | 16 | 10500 |
| P2 | 31 | 16148 | 18 | 9553 | 37 | 22384 | 19 | 13029 |
| P3 | 22 | 8363 | 9 | 3455 | 38 | 23286 | 13 | 7392 |
| P4 | 19 | 11322 | 15 | 9321 | 33 | 19459 | 14 | 9198 |
| P5 | 32 | 24376 | 16 | 12886 | 34 | 24290 | 15 | 9996 |

P1-P5: Participant 1-5; VL: vastus lateralis muscle, VM: vastus medialis muscle;

### 3.2. Generalizability of the deep CNN within participant

For the within-participant validation, the correlation coefficient was 0.983 ±0.004 (mean±std across 5 participants) for 3-fold CV and 0.977±0.007 for LOOCV (Fig. 4(a)). The nRMSE was 5.45±0.55% for 3-fold CV and 6.28±0.69% for LOOCV (Fig. 4(b)). Figure 5 shows that the neural drive from BSS and that from deep CNN are highly correlated (r=0.98±0.01; nRMSE=5.6±1.3) for all four muscle-session combinations from one representative participant in the LOOCV. Regardless of the validation method, the deep CNN trained and validated with data from P1 generated the best performance with a correlation coefficient of 0.982±0.005 and a nRMSE of 5.58±0.81%; when trained with data from P3, the deep CNN performed the worst resulting in a correlation coefficient of 0.967±0.008 and a nRMSE of 7.19±0.98%. For the correlation coefficient, the ANOVA detected no significant difference between data sources (p=0.22), no significant difference between validation methods (p=0.27), and no significant interaction between validation methods and data sources (p=0.99). For nRMSE, the ANOVA detected significant differences between data sources (p=0.04), significant differences between validation methods (p=0.01), but no significant interaction between validation methods and data sources (p=0.92).

**Figure 4.**
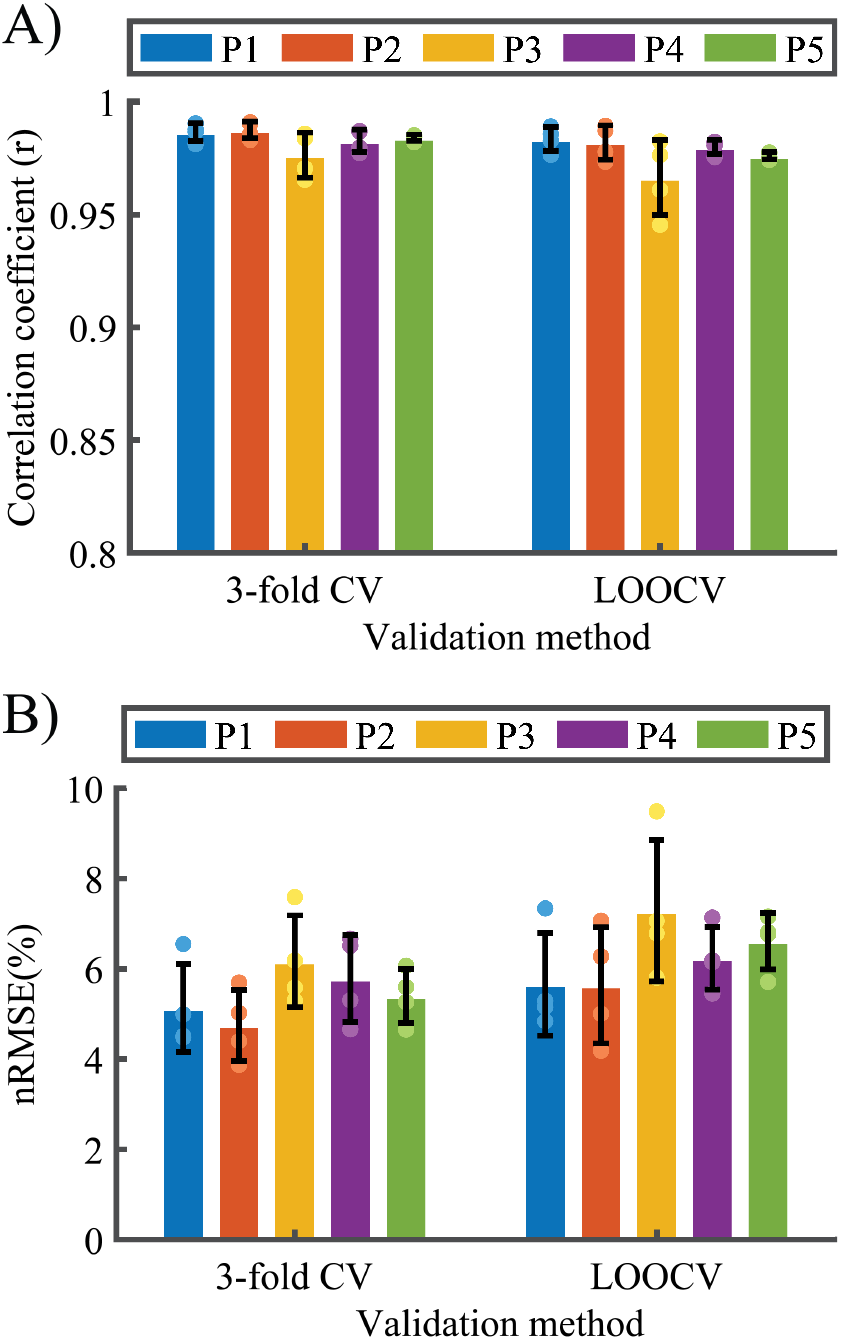
(A) The correlation coefficient between results of the BSS and deep CNN, (B) the nRMSE between results of the BSS and deep CNN from the within-participant cross validation. The validation performed using both 3-fold CV and LOOCV for five participants. The error bar represents the standard deviation from multiple runs.

**Figure 5.**
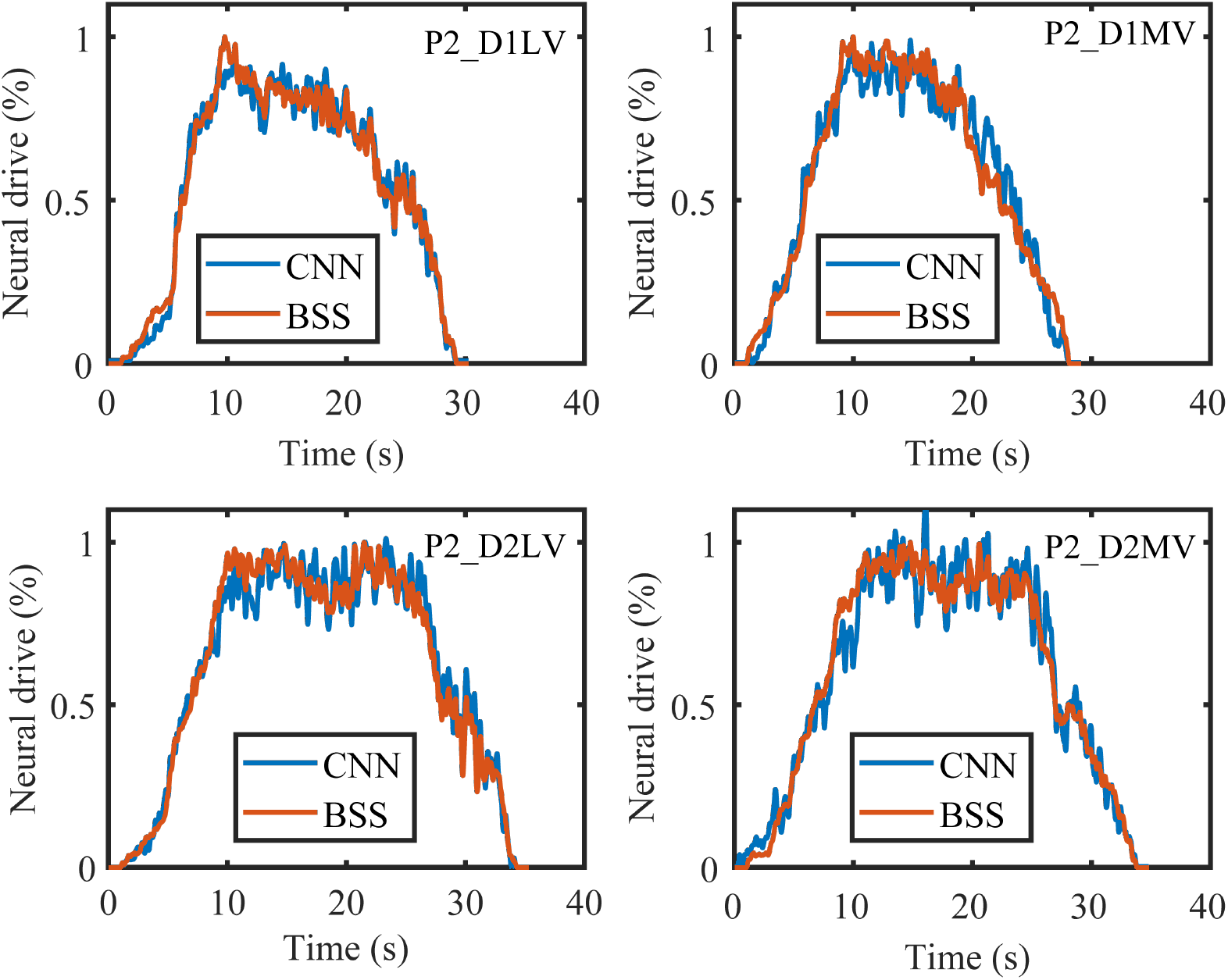
Comparison between the neural drive estimated using BSS and the neural drive estimated using the deep CNN for within-participant LOOCV. The deep CNN was trained with three muscle-session conditions and validated with the remaining one. Here are the results from participant 2.

### 3.3. Generalizability of the deep CNN across participants

For across-participant validation, the correlation coefficient was 0.983±0.005 (mean±std across 4 muscle-session combinations) for 3-fold CV and 0.985±0.005 for LOOCV (Fig. 6(a)). The nRMSE was 5.55± 0.69% for 3-fold CV and 5.12±0.68% for LOOCV (Fig. 6(b)). Regardless of the validation method, the deep CNN performed the best when trained and validated with data from S1L with a resulting correlation coefficient of 0.987±0.003 (mean±std across two validation methods) and nRMSE of 5.05±0.62% (Fig. 6). The deep CNN performed the worst when trained and validated with data from S1M. The resulting correlation coefficient was 0.975±0.009 and nRMSE was 6.43±1.18%. For both correlation coefficient and nRMSE, the ANOVA detected a significant difference between data sources (p*<*0.005), no significant difference between validation methods (p*>*0.12), and no significant interaction between validation methods and data sources (p=0.99).

**Figure 6.**
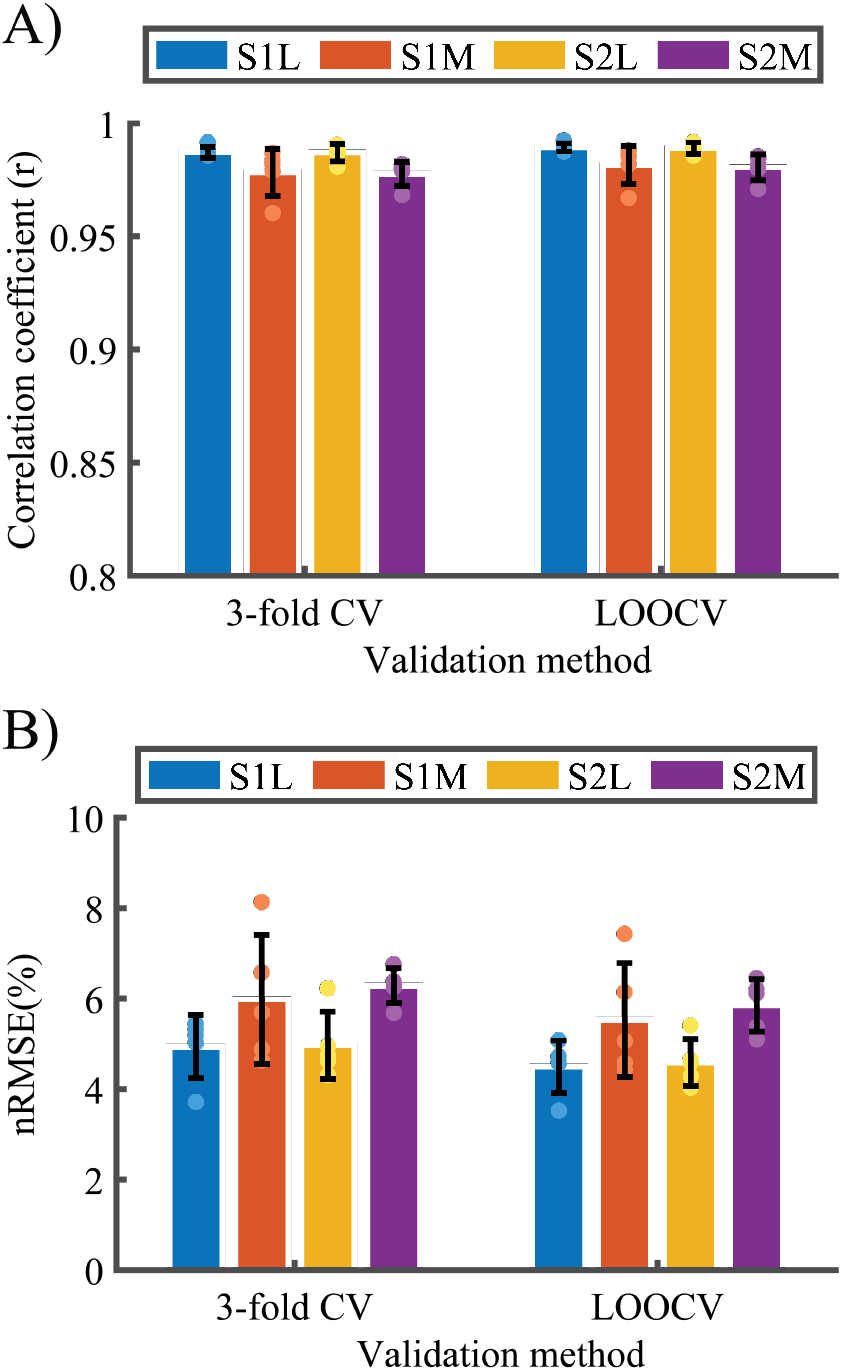
The correlation coefficient (A) and the nRMSE (B) of the deep CNN from the across-participant cross validation. The validation was performed using both 3-fold CV and LOOCV for four muscle-session conditions. The error bar represents the standard deviation from multiple runs.

### 3.4. Generalizability of the deep CNN across all conditions

The generalizability of the deep CNN across all conditions increased as more datasets were included in the training data (Fig. 7). Trained with only one dataset, the correlation coefficient was 0.97±0.007 (mean and standard deviation) across 20 runs, ranging from 0.953 to 0.977 (MIX1 in Fig. 7). Trained with four datasets, the correlation coefficient was 0.979±0.001 (mean and standard deviation) across 5 runs, ranging from 0.978 to 0.98 (MIX4 in Fig. 7). As shown in Fig. 8, the number of MUs in the training data has a significant effect on the average correlation coefficient across all testing conditions (r = 0.60 and p*<*0.01).

**Figure 7.**
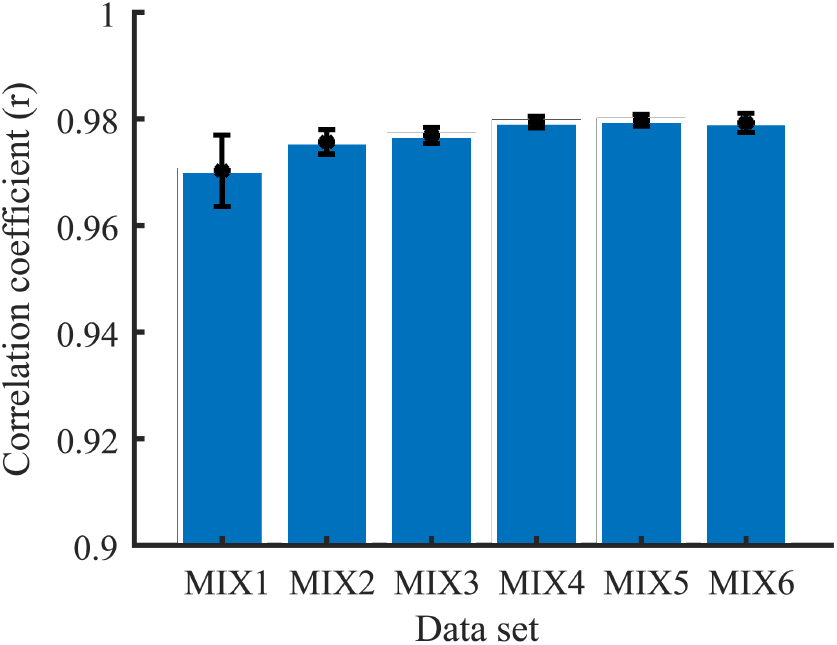
The generalizability of deep CNN across all training conditions with different numbers of datasets. The error bar represents the standard deviation across multiple runs in each condition.

**Figure 8.**
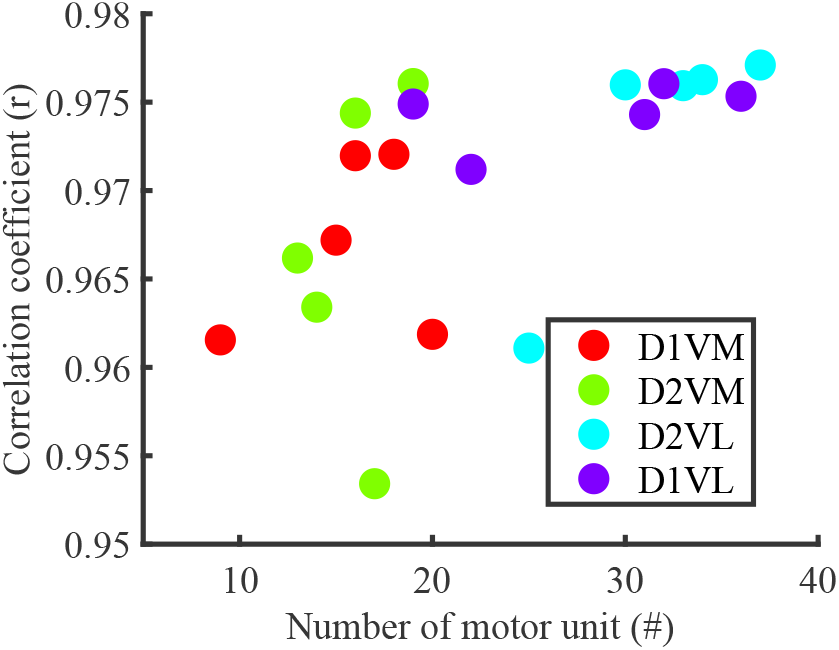
Relationship between the generalizability of deep CNN and the number of MUs in the training data. The training data included only one dataset (i.e. MIX1). There are 20 data points in total, including 5 participants each with 4 muscle-session conditions.

## 4. Discussion

This study aimed to validate the feasibility of using deep CNN for neural drive estimation from HD-EMG and to investigate the generalizability of this approach across different muscles, sessions, and participants. In principle, the deep CNN can learn the general features of MUAPs from a pool of MUs and estimate the neural drive from HD-EMG in an online manner. We recorded HD-EMG signals from the VL and VM muscles during two different sessions and decomposed the HD-EMG signals into corresponding MU spike trains using a blind source separation method. Then, the HD-EMG signals and the CST (summation of all MU spike trains) were segmented into data frames for the training and validation of the deep CNN. We performed 3-fold CV and LOOCV for within-participant and across-participant situations. Without retraining, the deep CNN is generalizable across different muscles and participants for long-term neural drive estimation. More important, in LOOCV, the deep CNN achieved similar performance as in 3-fold CV, indicating it is generalizable to new dataset with comparable performance. With a higher number of MUs included in the training data, the deep CNN is more generalizable to different muscles, sessions, and participants.

For data analysis, the ramp-up and ramp-down segments were included to take into account the performance in estimating the neural drive during linear changes in the force generation profile, which is typical in evaluating neural-machine interface [15–17]. The 2-s period of 0% MVC (i.e., non-active period) was used to account for false positives in estimating the neural drive. This is a trade-off to assess the false positives without inflation of the correlation coefficient with a long non-active period. The deep CNN performed reasonably well compared to a previous study on intra-muscular EMG decomposition, which reported correlation coefficients of 0.95±0.04 between estimated CST using an online method and offline manually edited CST using EMGlab decomposition software [38]. A few studies first estimated the neural drive using online methods and then used a linear regression model to estimate joint force using neural drive, and they reported correlation coefficients ranging from 0.83 to 0.91 [36, 39]. We reference these studies but cannot make fair comparison between our results and theirs because of differences in selection of muscles and experimental setups.

The performance of the deep CNN varied slightly for each dataset (e.g., a lower correlation coefficient for participant 3) in Fig. 4 and Fig. 6. We speculate that the performance of the deep CNN was related to the number of MUs and the number of spikes in the training data. In Table. 1, we observed that 1) participant 3 has fewer MUs and spikes than other participants and 2) the VM muscle has fewer MUs and spikes than VL muscle for both sessions across all participants. These observations are aligned with the performance decreases with certain datasets. This is also supported by the generalizability validation of the deep CNN across all conditions. The number of MUs was significantly positively correlated with the averaged correlation coefficient across all testing datasets (Fig. 8). To make the results more generalizable, we will train the deep CNN with benchmark HD-EMG data and validate with our experimental data in our future study.

In this study, we performed a few pre-processing procedures, such as visual inspection of the HD-EMG signals to remove noisy channels, that need to be automated for online application. Current efforts have been made to automate removal of noisy channels using techniques such as independent component analysis [40], spatial similarity via normalised mutual information between electrodes [41], and outlier detection via local distance-based outlier factor [42]. To this end, we could easily integrate one of these methods to our framework or include a calibration step to remove noisy channels. After automatic removing of noisy channels, we will investigate the robustness of the deep CNN method and how the number of missing channels affects its performance, as this will determine to what extend the deep CNN can reliably estimate the neural drive in real applications.

Additionally, we have only investigated the generalization across the VL and VM muscles. VL and VM muscles have very similar functions in terms of actuating the knee, leading to a high cross-correlation between the two muscle activities. This might have led to the promising results in the generalization between the two muscles. In our future work, we will test the generalization across muscles with higher independence and greater anatomical differences (e.g., generalization between the tibialis anterior muscle and the gastrocnemius muscle). Currently, MU identification requires manual-edits by an operator, which is time consuming and labor intensive. If simulated data (i.e., simulated HD-EMG signals) can be used to train a deep CNN that can be generalizable for different muscles, this can greatly improve the practicality of this approach to reduce the calibration process.

## 5. Conclusion

We proposed a deep CNN framework to estimate the neural drive in the form of CST from HD-EMG signals across different muscles, sessions, and participants. Our study demonstrated that 1) the deep CNN is generalizable to different muscles and participants, i.e., estimating the neural drive for conditions that are not included in the training data, and 2) the deep CNN is scalable to capture complex features of variant MUAP shapes when trained with combined data that are recorded from different muscles and participants. Compared with the commonly used offline neural drive extraction approach (i.e., BSS), the proposed deep CNN framework could accurately estimate the neural drive using HD-EMG signals. Moreover, the proposed deep CNN is generalizable to different muscles and participants without retraining and could run in a realtime manner as was done in a previous study [32]. Therefore, the proposed deep CNN framework is a promising candidate to extract neural drive for realtime human-machine interface for assistive technology (e.g., exoskeletons and prostheses).

